# Linear Double-stranded DNAs as Innovative Biological Parts to Implement Genetic Circuits in Mammalian Cells

**DOI:** 10.1101/266056

**Authors:** Shuai Li, Weijun Su, Chunze Zhang

**Affiliations:** Department of Breast Cancer Pathology and Research Laboratory, Tianjin Medical University Cancer Institute and Hospital, National Clinical Research Center for Cancer, Key Laboratory of Cancer Prevention and Therapy, Tianjin 300060, China, Tianjin’s Clinical Research Center for Cancer; School of Medicine, Nankai University, Tianjin 300071, China; Department of Colorectal Surgery, Tianjin Union Medical Center, Tianjin 300121, China

**Keywords:** synthetic biology, genetic circuit, Boolean AND gate, linear double-stranded DNA (ldsDNA), non-homologous end joining (NHEJ)

## Abstract

Synthetic biology employs engineering principles to re-design biological systems for biomedical or industrial purposes. Innovation and application of original biological parts for genetic circuit construction will significantly facilitate and expedite the development of synthetic biology. Here, we built two- or three-input linear double-stranded DNA (ldsDNA)-based Boolean AND gate genetic circuits in mammalian cells. Bioluminescence living imaging revealed the feasibility of ldsDNA-based Boolean AND gate circuits *in vivo*. Inhibition of DNA-PKcs, a pivotal enzyme in non-homologous end joining (NHEJ), significantly attenuated the output signals from ldsDNA-based Boolean AND gate circuits. We further showed that ldsDNA with terminal additional random nucleotide(s) could undergo end nucleotide deletion and generate in-frame protein via the Boolean AND gate response. Additionally, ldsDNAs or plasmids with identical overlapping sequences could also serve as input signals for Boolean AND gate genetic circuits. Our work establishes ldsDNA as innovative biological parts for building low noise-signal ratio Boolean AND gate circuits with application potential in biomedical engineering fields.

## Introduction

One primary focus of synthetic biology is on designing genetic circuits for biomedical or industrial purposes [1-3]. To date, a series of biological parts with different underlying compositions (DNA, RNA, or protein) have been developed to function as basic building blocks of genetic circuits. DNA is commonly used as biological parts, functioning as either a regulator or adaptor in genetic circuits [4]. For instance, synthetic transcription factors could bind to specific DNA sequences and turn on gene expression [5-9]. DNA recombinases can recognize recombination DNA sequences and catalyze recombination, thereby accomplishing sophisticated biocomputation [10, 11]. The innovation of new DNA-based biological parts would help advance genetic circuit design.

Similar to complex electronic circuits, genetic circuits accomplish biocomputing by integrating different logic gates into networks. Among these logic gates, the Boolean AND gate converts two (or more) input signals into one output signal through Boolean calculation, which can be exploited to enhance diagnostic or therapeutic specificity in biomedical studies [12-15]. Over the past two decades, various Boolean AND logic gate design strategies have been developed. These strategies could be divided into two subtypes: ‘circuit in place’ and ‘bio-parts self-assembly’. For the commonly used ‘circuit in place’ strategy, input signal integration is executed in the presence of all computation modules. For the ‘bio-parts self-assembly’ Boolean AND-gate strategy, the TRUE output signal could be generated only when all inputs (or their products) are assembled as one output-signal-generating molecule [16-18].

Here, we used linear double-stranded DNA (ldsDNA, or ‘PCR amplicon’) as innovative biological parts to implement Boolean AND logic gates. Via splitting the essential gene expression cassette into signal-mute ldsDNAs, this strategy results in low noise-signal ratio Boolean AND gate circuits both *in vitro* and *in vivo*. Additionally, ldsDNA with one or two additional terminal nucleotide(s), which leads to a reading frame shift, could undergo end-processing and produce in-frame Boolean AND gate output proteins. Moreover, ldsDNAs or plasmids with identical overlapping sequences could also implement Boolean AND gate circuits. Our present study provides novel biological parts and principles for synthetic genetic circuit design, which may have application potential in biomedical engineering fields.

## Results

### Two-input ldsDNA-based Boolean AND gate genetic circuits in mammalian cells

A typical gene overexpression plasmid (for mammalian cells) contains an integrant sequence consisting of a promoter, gene coding sequence (CDS) and poly(A) signal (PAS). Here, by PCR amplification, we split the reporter gene expression cassette (promoter-CDS-PAS) into 2 or 3 ldsDNAs. Then, these input ldsDNAs were introduced into mammalian cells to construct Boolean AND gate genetic circuits.

Taking the pcDNA3.0-firefly luciferase expression vector as the PCR template, we first designed three different pairs of two-input ldsDNA-based Boolean AND gate genetic circuits (Figure 1A). ldsDNAs were synthesized by PCR amplification, and the agarose gel electrophoresis results are shown in Figure 1B. State [0,0], [0,1], [1,0], or [1,1] Boolean AND gate genetic circuits were separately introduced into HEK293T cells by transfection. To balance the transfection efficiency, we cotransfected the CMV-Renilla luciferase plasmid (pRL-CMV). As shown in Figure 1C, state [1,1] led to a dramatic induction of relative luciferase activity (∼16-, 7-, and 26-thousand times in each pair) compared to that of non-ldsDNA transfected cells (state [0,0]). A similar response was achieved with genetic circuits constructed with *de novo* synthesized commercialized DNA (data not shown). A similar trend was observed when firefly luciferase activities were normalized to protein concentrations (Figure 1D). Raw firefly luciferase relative luminescence unit (RLU) values are presented in Figure 1E. However, as shown in Figure 1F, the luciferase reporter levels generated by this system were still lower than that from ldsDNA with an intact gene-expression cassette (CMV-Luc[1-1653]-BGH PAS). There was a positive correlation between the amount of input ldsDNA and the intensity of output signals (Figure 1G).

**Figure 1.**
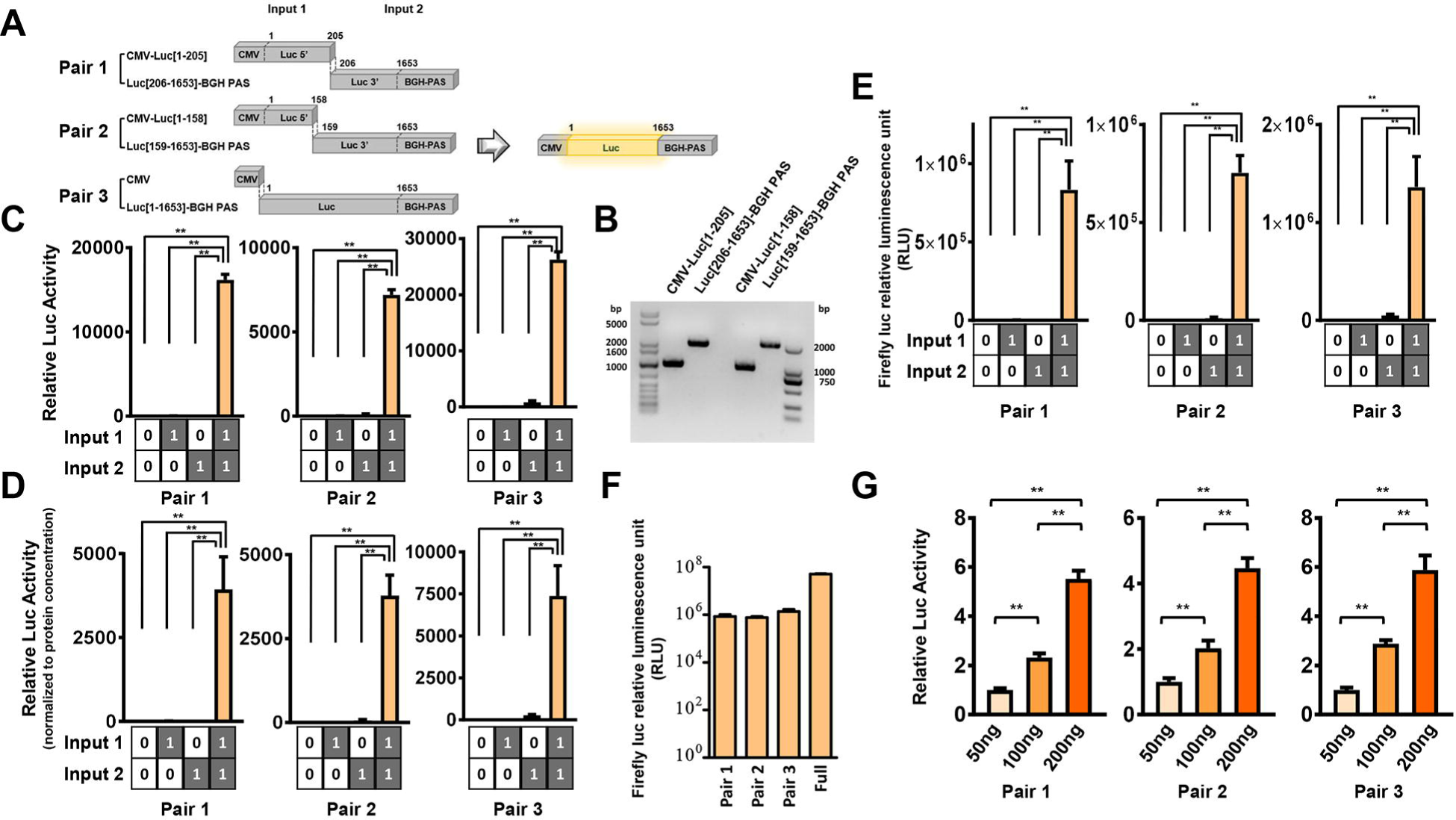
Induction of firefly luciferase expression via two-input ldsDNA-based Boolean AND gate circuits *in vitro*. (A) Schematic of 3 pairs of pcDNA3.0-firefly luciferase vector derived two-input ldsDNA-based Boolean AND gate. 1^st^ pair: input 1 as CMV-Luc[1-205], input 2 as Luc[206-1653]-BGH poly(A) signal (PAS); 2^nd^ pair: input 1 as CMV-Luc[1-158], input 2 as Luc[159-1653]-BGH PAS; 3^rd^ pair: input 1 as CMV, input 2 as Luc[1-1653]-BGH PAS. (B) Agarose gel electrophoresis of ldsDNA PCR products (CMV-Luc[1-205], Luc[206-1653]-BGH PAS, CMV-Luc[1-158] and Luc[159-1653]-BGH PAS). (C-F) The indicated two-input ldsDNA-based genetic circuits were introduced into HEK293T cells for 48 h. (C) Firefly luciferase activity was normalized to cotransfected Renilla signals. (n = 4). (D) Firefly luciferase activity was normalized to protein concentrations. (n = 4). (E) Firefly luciferase output signals of states [0,0], [1,0], [0,1] and [1,1] were measured and raw RLU values are shown. (n = 4). (F) Firefly luciferase output signals from different Boolean AND gate states [1,1] or ldsDNA with intact gene expression cassette (full, CMV-luc[1-1653]-BGH PAS) were measured and the RLU values are shown. (n = 4). (G) Different amounts of ldsDNA were transfected into HEK293T cells for 48 h as indicated. Firefly luciferase output signals were measured and normalized to cotransfected Renilla signals. (n = 4). All data are displayed as the mean ± SD; \*\**p* < 0.01; statistical significance calculated using two-tailed Student’s t-test.

Next, we designed a two-input ldsDNA-based Boolean AND gate genetic circuit by amplifying the CMV promoter fragment and the GFP coding sequence-SV40 poly(A) signal fragment (GFP-SV40 PAS) from the pEGFP-C1 vector (Figure 2A). Flow cytometry showed that the Boolean AND gate response led to a 20.5% GFP-positive rate in transfected cells (state [1,1]) (Figure 2B-C). Moreover, western blot results also demonstrated the expression of GFP protein with the introduction of state [1,1] (Figure 2D).

**Figure 2.**
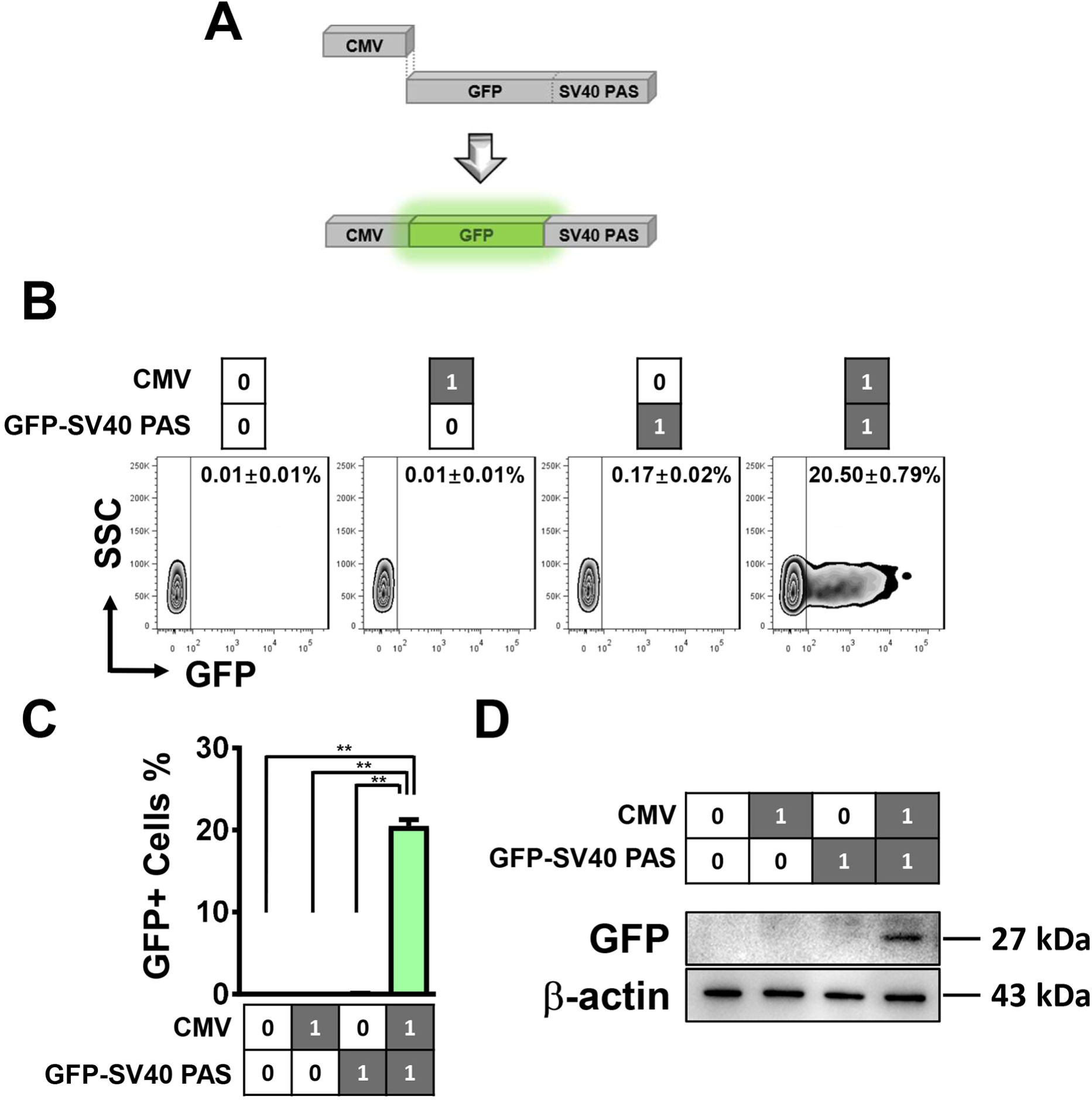
Induction of GFP expression via two-input ldsDNA-based Boolean AND gate circuits *in vitro*. (A) Schematic of the pEGFP-C1 vector-derived two-input ldsDNA-based Boolean AND gate. Input 1 as the CMV promoter, input 2 as GFP-SV40 PAS. (B) The indicated two-input ldsDNA-based genetic circuits were transfected into HEK293T cells for 48 h, and GFP-positive rates of states [0,0], [1,0], [0,1] and [1,1] were analyzed by FACS. (C) Statistics of FACS data in (B). (n = 3). (D) Western blot analysis of GFP expression under the indicated states. Data are displayed as the mean ± SD; \*\**p* < 0.01; statistical significance calculated using two-tailed Student’s t-test.

In the experimental plasmid ligation or cellular non-homologous double-strand end joining (NHEJ) process during double-strand DNA break repair, a covalent phosphodiester bond is formed between the 5′ phosphate group from one dsDNA and the 3′ hydroxyl group from another dsDNA [19, 20]. As we know, regular synthesized PCR primers contain a 5′ terminal hydroxyl group (not a phosphate group), which leads to hydroxyl groups present on both ends of the PCR amplicons. Here, with 5′ phosphorylated primers, we generated 5′ terminal phosphorylated ldsDNA by PCR (taking the pcDNA3.0-firefly luciferase plasmid as a template). However, unexpectedly, no significant enhancement of reporter activities was observed in 5′ phosphorylated ldsDNA-based Boolean AND genetic circuits compared to non-phosphorylated ldsDNAs (Figure 1C and Figure 3).

**Figure 3.**
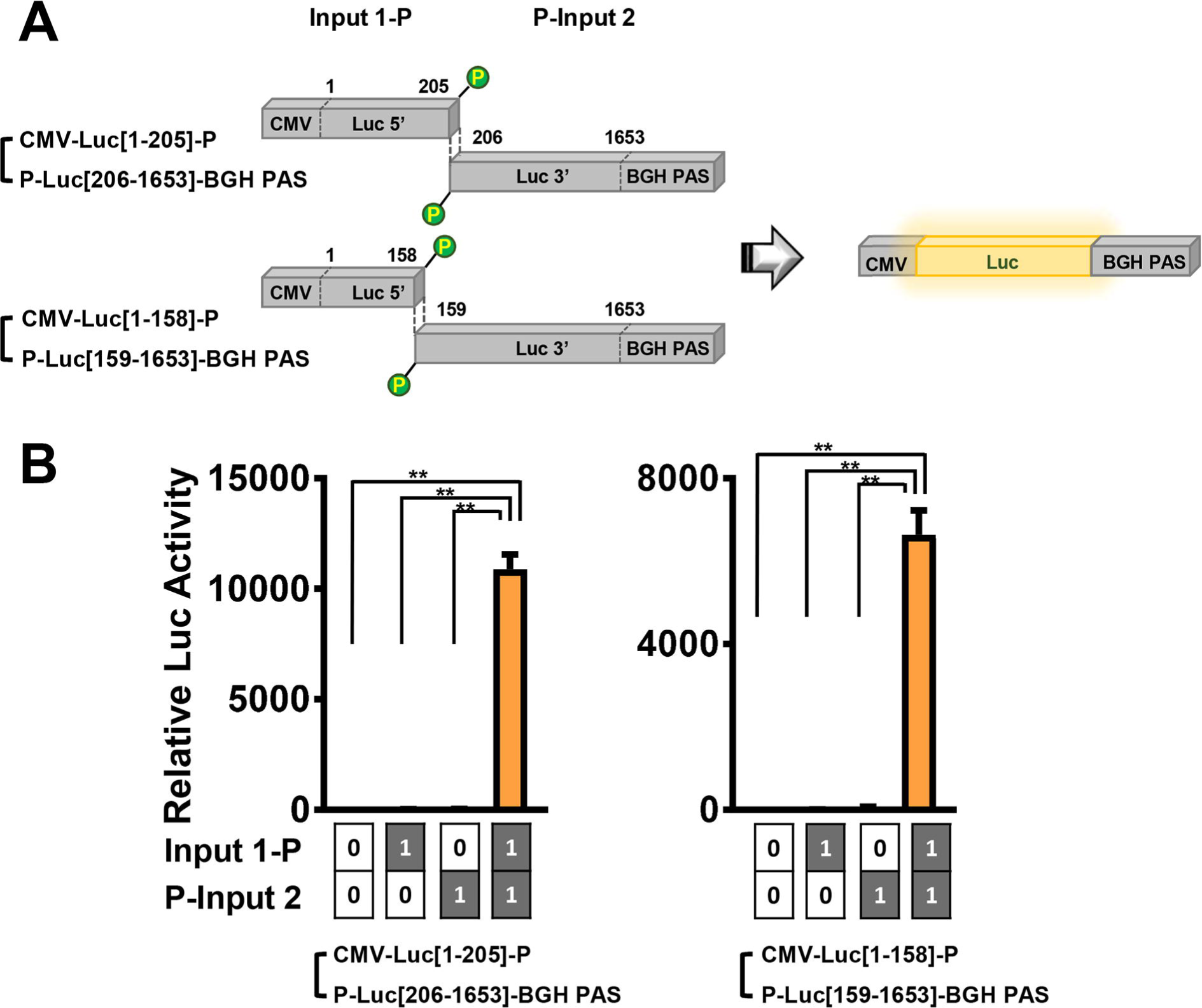
Two-input Boolean AND gate circuits with 5′ terminal phosphorylated ldsDNA. (A) Schematic of 2 pairs of two-input Boolean AND gate 5′ terminal phosphorylated ldsDNA amplified from the pcDNA3.0-firefly luciferase plasmid. 1^st^ pair: input 1-P as CMV-Luc[1-205]-P, P-input 2 as P-Luc[206-1653]-BGH PAS; 2^nd^ pair: input 1-P as CMV-Luc[1-158]-P, P-input 2 as P-Luc[159-1653]-BGH PAS. (B) The indicated 5′ terminal phosphorylated ldsDNA-based genetic circuits were transfected into HEK293T cells, and the firefly luciferase output signals of states [0,0], [1,0], [0,1] and [1,1] were measured and normalized to cotransfected Renilla signals. (n = 4). The experiments were performed simultaneously with that in Figure 1C. All data are displayed as the mean ± SD; \*\**p* < 0.01; statistical significance calculated using two-tailed Student’s t-test.

### Inhibition of DNA-PKcs attenuates output signals of ldsDNA-based Boolean AND genetic circuits

Because the ldsDNA end-relinkage process is similar to the NHEJ pathway, we further explored whether inhibition of the NHEJ pathway could attenuate Boolean AND gate output signals. DNA-PKcs (DNA-dependent protein kinase, catalytic subunit), a pivotal Ku70/Ku80-interacting nuclear serine/threonine kinase, was selected to be targeted by RNA interference (Figure 4A) [21]. As shown in Figure 4B, the inhibition of DNA-PKcs significantly downregulated ldsDNA-based Boolean AND gate output signals.

**Figure 4.**
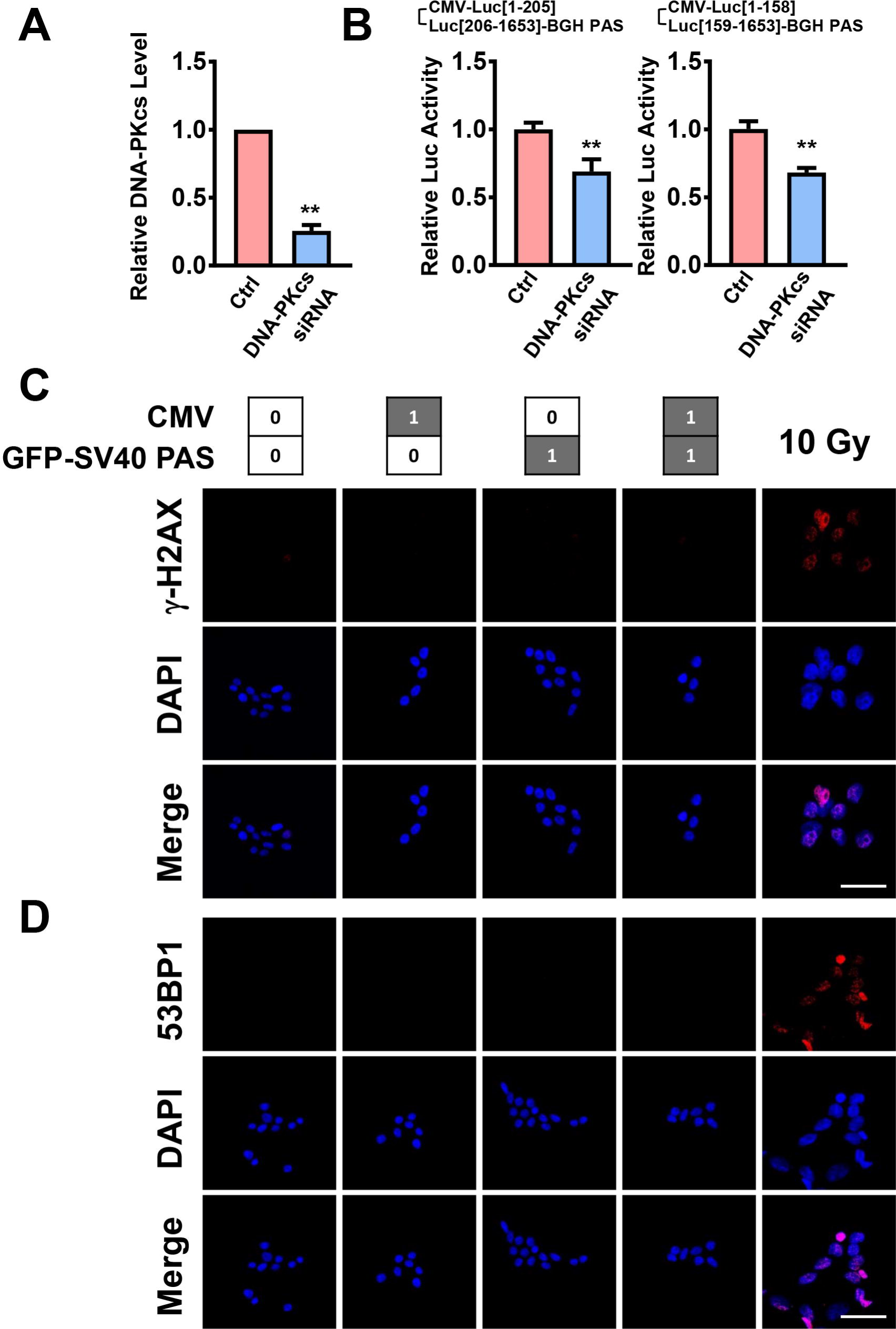
Attenuation of ldsDNA-based Boolean AND gate circuit output signals by DNA-PKcs siRNA and detection of DNA damage foci formation in cells transfected with ldsDNA. (A) qRT-PCR analysis of the DNA-PKcs siRNA effect in HEK293T cells. (n = 3). (B) HEK293T cells were transfected with ldsDNA-based Boolean AND gate genetic circuits as indicated. Relative firefly luciferase output activities (normalized to cotransfected Renilla signals) of the indicated Boolean AND gate circuits (state [1,1]) in DNA-PKcs knockdown cells are shown here. (n = 4). (C-D) HEK293T cells were transfected with ldsDNA-based Boolean AND gate genetic circuits as indicated. After 48 h, immunofluorescence staining of the DNA damage foci markers γ-H2AX and 53BP1 was performed. Cells undergoing 10 Gy γ-radiation served as positive controls. Representative images of γ-H2AX and 53BP1 staining are shown in (C) and (D), respectively. Scale bar = 50 μm. All data are displayed as the mean ± SD; \*\**p* < 0.01; statistical significance calculated using two-tailed Student’s t-test.

### Introduction of ldsDNA does not lead to DNA damage foci formation

Given the potential applications of this system in biomedical treatments, we investigated whether ldsDNA introduction could lead to DNA damage foci formation in cells. After ldsDNA transfection, immunofluorescence staining of DNA damage foci markers, such as γ-H2AX and 53BP1, was performed in HEK293T cells, and 10 Gy γ-radiated cells were used as a positive control. As shown in Figure 4C-D, neither γ-H2AX nor 53BP1 foci formed in ldsDNA-transfected cells.

### ldsDNA-based two-input Boolean AND gate genetic circuits *in vivo*

To estimate whether ldsDNA-based Boolean AND gate genetic circuits could be constructed *in vivo*, we employed hydrodynamic injection as a tool for *in vivo* ldsDNA delivery [22]. First, we confirmed the validity of this method by rapidly injecting a large volume of CMV-Luc[1-1653]-BGH PAS ldsDNA solution (full gene expression cassette) into the mouse retro-orbital sinus. With bioluminescence imaging (BLI), firefly luciferase signals were detected in the liver region one day after hydrodynamic injection and diminished in 2 weeks (Figure 5A-B). Postmortem BLI confirmed the expression of luciferase in the liver but no other organs (Figure 5C), which is consistent with the results from other groups [22]. We next introduced state [0,1], [1,0] or [1,1] Boolean AND gate genetic circuits into mice. State [1,1] (CMV-Luc[1-205] + Luc[206-1653]-BGH PAS), but not state [0,1] or [1,0], led to firefly luciferase expression *in vivo* (Figure 5D-E). Correspondingly, a significant luciferase signal was detected in the liver under state [1,1] via postmortem BLI (Figure 5F-G).

**Figure 5.**
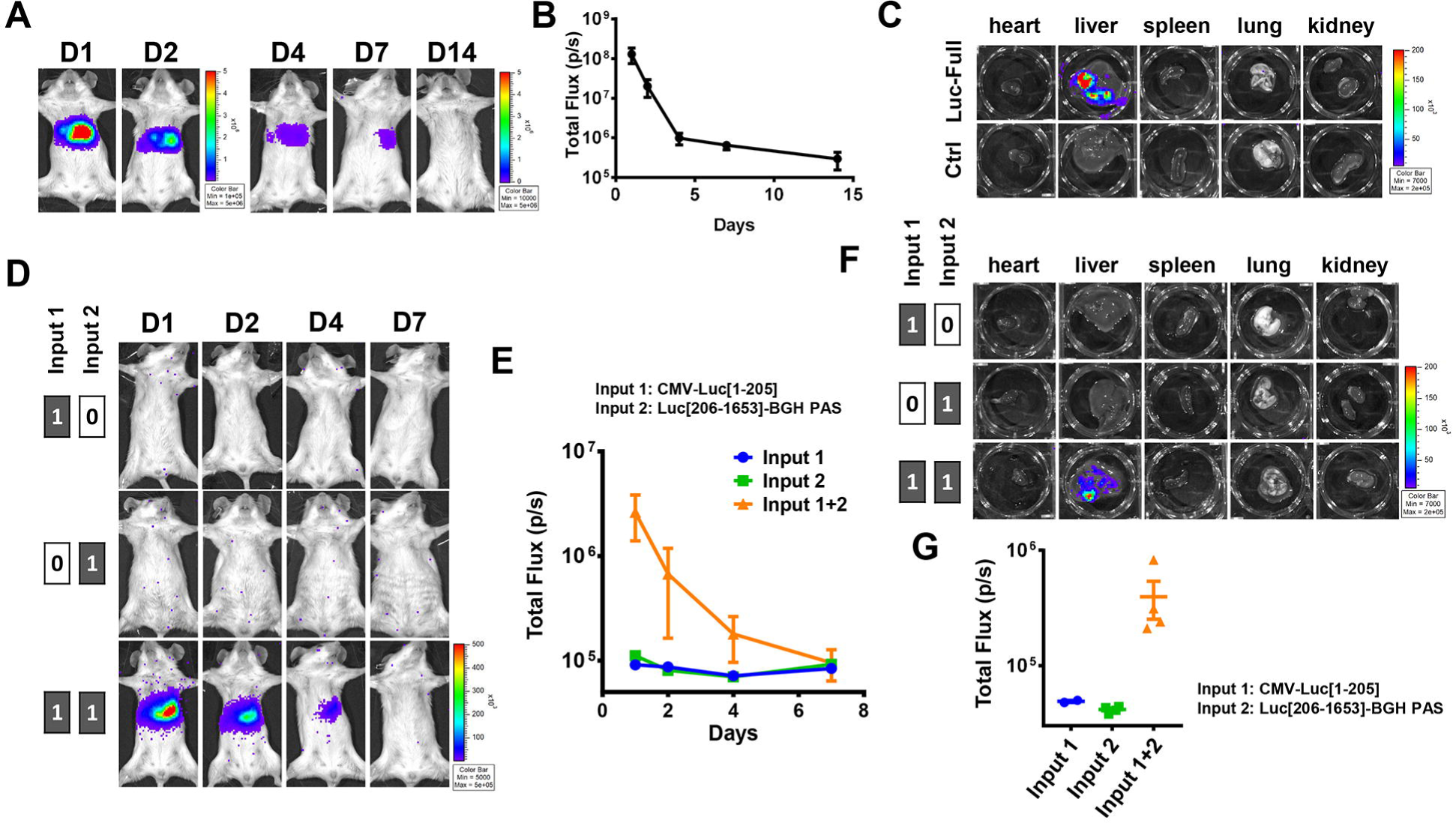
Two-input ldsDNA-based Boolean AND gate circuits *in vivo*. (A-C) Bioluminescence imaging (BLI) of firefly luciferase in mice undergoing CMV-Luc[1-1653]-BGH PAS ldsDNA hydrodynamic injection. Representative images at different time points are shown in (A). Signals were quantified in units of maximum photons per second (photons/s), and the statistics are shown in (B). Data are displayed as the mean ± SEM. (n = 4). (C) Mice were euthanized one day after hydrodynamic injections, and firefly luciferase BLI was performed with different organs. Representative images are shown here. (D-G) The indicated ldsDNA inputs were introduced into mice via hydrodynamic injection through the retro-orbital sinus, and firefly luciferase BLI was performed at the indicated time points. Representative images of the indicated time points are shown in (D). Signals were quantified in units of maximum photons per second (photons/s), and the statistics are presented in (E). Data are displayed as the mean ± SEM. (input 1: n = 5, input 2: n = 4, input 1 + 2: n = 5). (F-G) Mice were euthanized one day after hydrodynamic injections of ldsDNA, and firefly luciferase BLI was performed with different organs. Representative images are shown in (F). Signals of the liver were quantified in units of maximum photons per second (photons/s), and the statistics are presented in (G). Data are displayed as the mean ± SEM. (input 1: n = 2, input 2: n = 3, input 1 + 2: n = 4).

### Three-input ldsDNA-based Boolean AND gate genetic circuits in mammalian cells

We further explored the possibility of building three-input ldsDNA-based Boolean AND logic circuits in mammalian cells. Three ldsDNAs (CMV promoter, GFP coding sequence, and SV40 PAS) were amplified from the pEGFP-C1 plasmid and transfected into HEK293T cells with different combinations (Figure 6A). Approximately 8% of the cells expressed GFP when all three ldsDNAs were introduced (state [1,1,1]) (Figure 6B-C). This finding implies that an increase in the number of inputs may lead to a decrease in the GFP-positive rates (two-input vs. three-input, 20.5% vs. 8.01%, Figures 2C and 6C).

**Figure 6.**
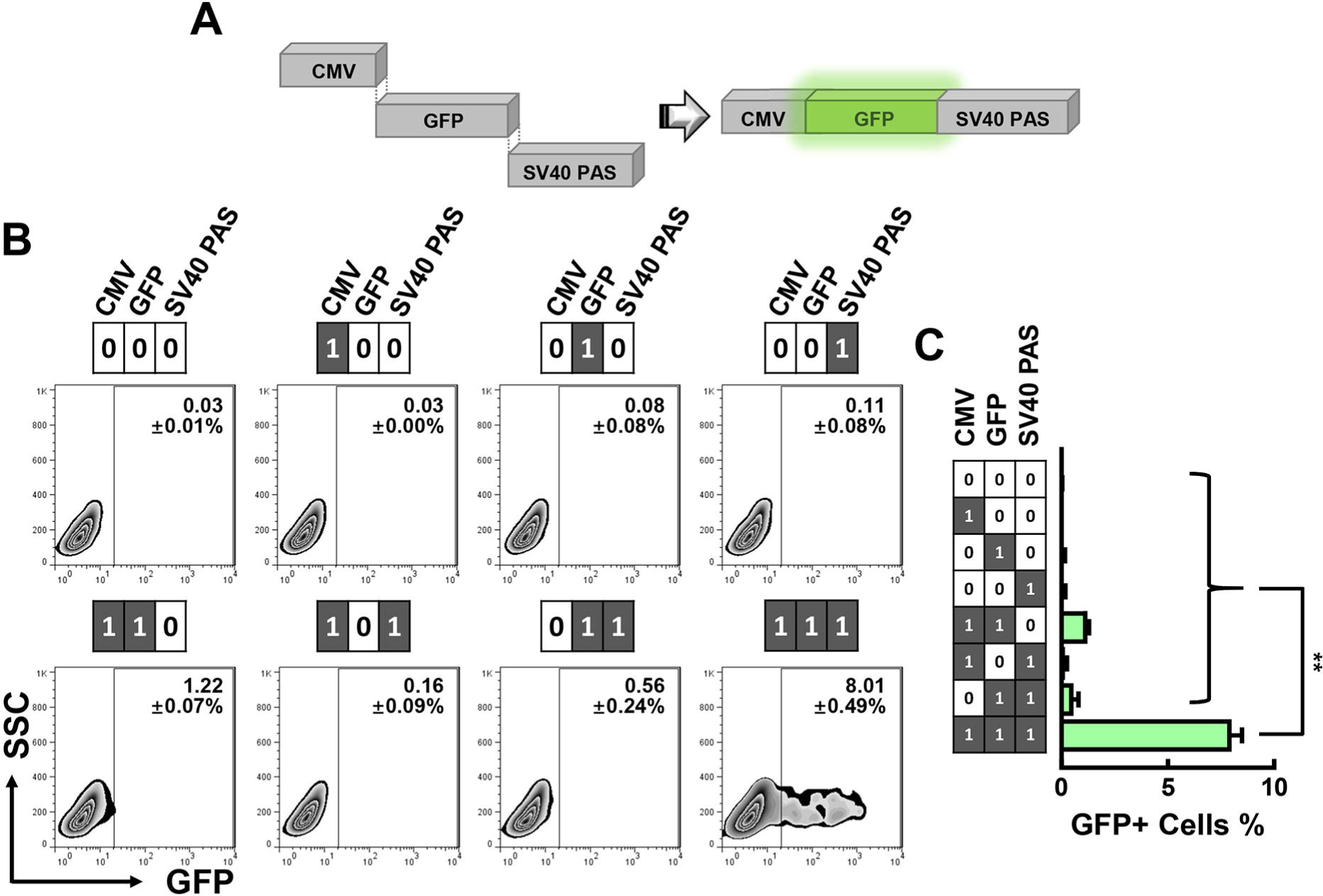
Three-input ldsDNA-based Boolean AND gate circuit. (A) Schematic of the three-input ldsDNA-based Boolean AND gate. Input 1 as the CMV promoter, input 2 as GFP CDS, and input 3 as SV40-PAS. (B) The indicated three-input ldsDNAs were transfected into HEK293T cells for 48 h, and the GFP-positive rates were analyzed by FACS. (C) Statistics of FACS data in (B). (n = 3). Data are displayed as the mean ± SD; \*\**p* < 0.01; statistical significance calculated using two-tailed Student’s t-test.

### Boolean AND genetic circuits with ldsDNA supplemented with terminal additional nucleotide(s)

In NHEJ-mediated DNA damage repair, dsDNA ends undergo end processing, during which nucleotide insertion or deletion occurs at DNA terminals. Thus, we wondered whether introduced exogenous ldsDNA was also subject to similar DNA end processing. By PCR amplification, we added one or two random nucleotide(s) to PCR amplicon terminals (Figure 7A). Theoretically, no in-frame GFP protein would be produced without DNA end processing. However, in practice, pEGFP-C1 vector-derived ldsDNA supplemented with one or two additional terminal nucleotide(s) led to a 10.76% or 6.77% GFP-positive rate, respectively (Figure 7B-C). These results demonstrated that ldsDNA could undergo end nucleotide deletion and thereby produce in-frame proteins via Boolean AND genetic circuits.

**Figure 7.**
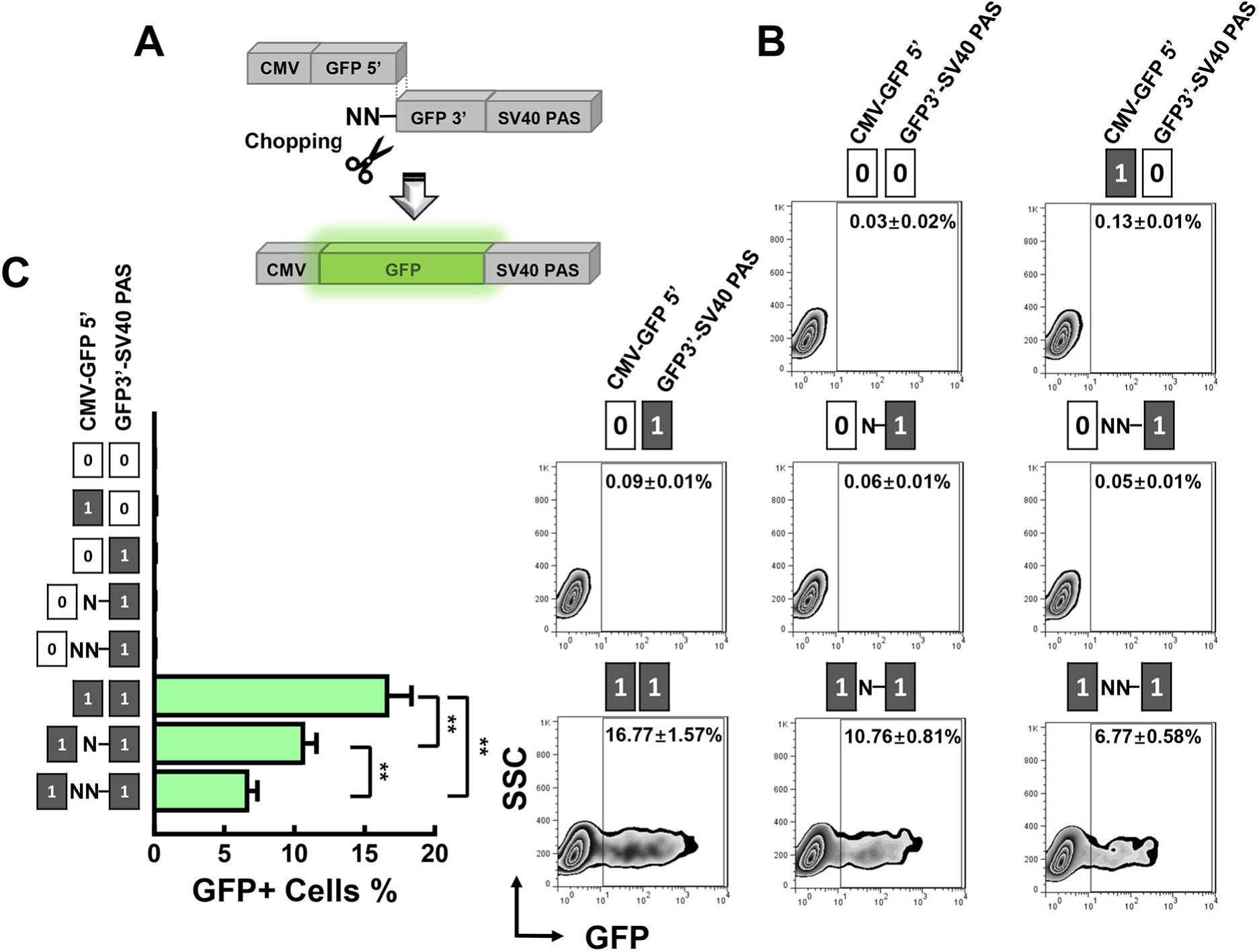
Boolean AND gate circuits by ldsDNA with random terminal additional nucleotide(s). (A) Schematic of two-input Boolean AND gate circuits using ldsDNAs with terminal additional nucleotide(s). Input 1 as CMV-GFP[1-204], input 2 as GFP[205-717]-SV40 PAS, N-GFP[205-717]-SV40 PAS or NN-GFP[205-717]-SV40 PAS. (B) The indicated two-input ldsDNAs with or without terminal additional random nucleotide(s) were transfected into HEK293T cells for 48 h, and the GFP-positive rates under the indicated states were analyzed by FACS. (C) Statistics of FACS data in (B). (n = 3). Data are displayed as the mean ± SD; \*\**p* < 0.01; statistical significance calculated using two-tailed Student’s t-test.

### ldsDNAs with identical overlapping sequences can construct Boolean AND logic circuits

We further revealed that ldsDNAs with identical overlapping sequences could serve as inputs of Boolean AND logic circuits (Figure 8A-B). The dual luciferase assay showed that ldsDNA pairs with different lengths of identical overlapping sequences led to 56-, 73-, 77- or 63- thousand times increases in output signals compared to that of nontransfected cells (state [0,0]) (Figure 8B), which was stronger than that by ldsDNA end rejoining (Figure 1C VS 8B). However, we found that longer overlapping sequences did not necessarily lead to higher output signal intensity (Figure 8B), while there was a positive correlation between the amount of input ldsDNA and the output signal intensity (Figure 8C). RNA interference knockdown of Rad51, a protein in charge of homologous searching in homologous recombination (HR), could attenuate output signals under state [1,1] (Figure 8D-E) [23]. Additionally, flow cytometry results showed that sequence-overlapping ldsDNAs (PCR-amplified from the pEGFP-C1 vector) could lead to GFP expression in 34.63% of the transfected HEK293T cells (Figure 8F-H).

**Figure 8.**
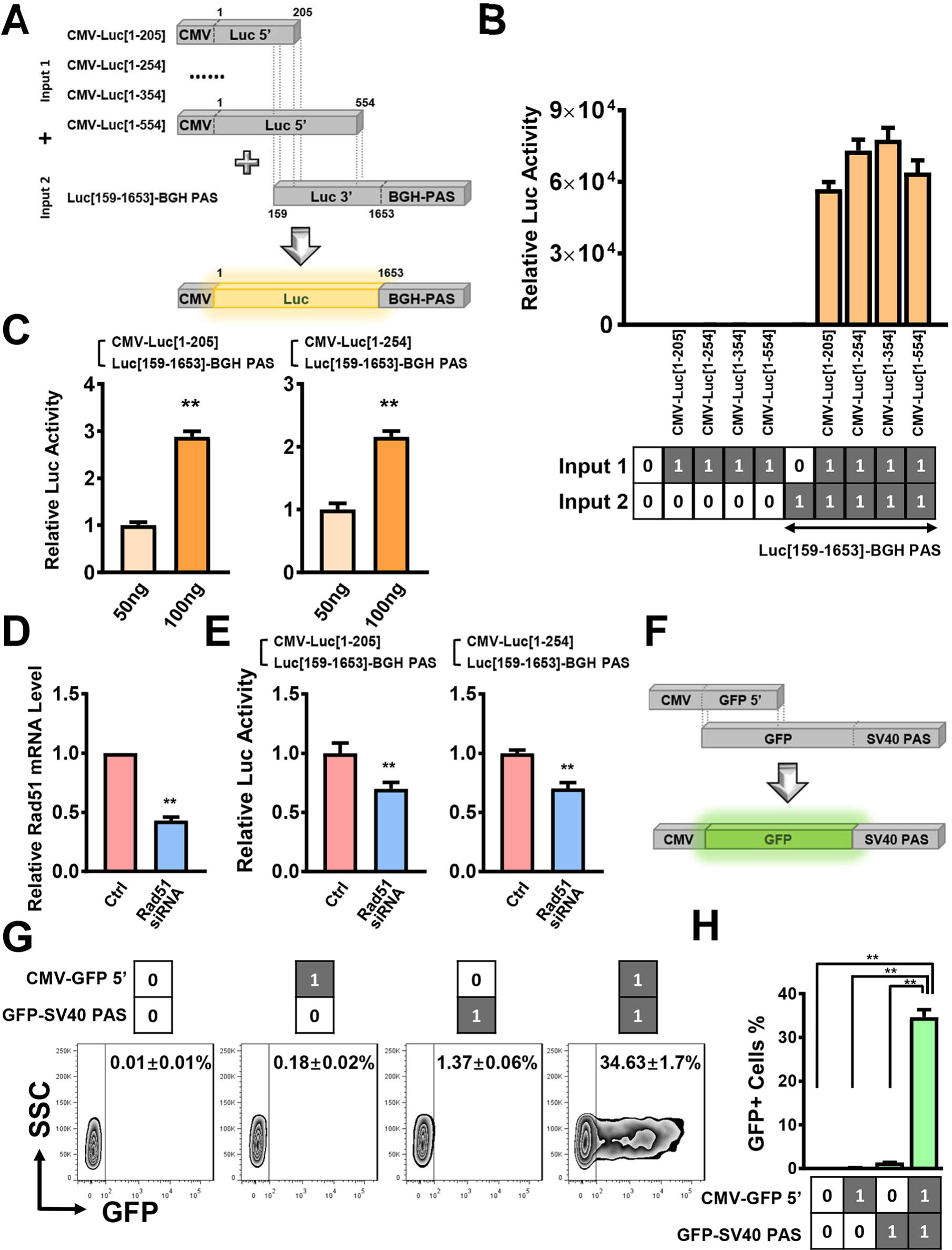
Boolean AND gate circuits by ldsDNAs with identical overlapping sequences. (A) Schematic of 4 pairs of pcDNA3.0-firefly luciferase vector-derived two-input Boolean AND gate by ldsDNAs with identical overlapping sequences. 1^st^ pair: input 1 as CMV-Luc[1-205], input 2 as Luc[159-1653]-BGH PAS; 2^nd^ pair: input 1 as CMV-Luc[1-254], input 2 as Luc[159-1653]-BGH PAS; 3^rd^ pair: input 1 as CMV-Luc[1-354], input 2 as Luc[159-1653]-BGH PAS; 4^th^ pair: input 1 as CMV-Luc[1-554], input 2 as Luc[159-1653]-BGH PAS. (B) The indicated two-input ldsDNA pairs were transfected into HEK293T cells for 48 h, and the firefly luciferase output signals were measured and normalized to cotransfected Renilla signals. The experiments were performed simultaneously with that in Figure 1C. (n = 4). (C) Different amounts of two-input ldsDNAs with identical overlapping sequences were transfected into HEK293T cells as indicated. Firefly luciferase output signals were measured and normalized to cotransfected Renilla signals. (n = 4). (D) qRT-PCR analysis of Rad51 siRNA knockdown efficiency in HEK293T cells. (n = 3). (E) Relative firefly luciferase output activities of the indicated Boolean AND gate circuits (state [1,1]) in Rad51 knockdown cells. (n = 4). (F) Schematic of the pEGFP-C1 vector-derived two-input Boolean AND gate by ldsDNAs with identical overlapping sequences. Input 1 as CMV-GFP[1-187], input 2 as GFP[1-717]-SV40 PAS. (G) The indicated ldsDNA pair was transfected into HEK293T cells for 48 h, and the GFP-positive rates of states [0,0], [1,0], [0,1] and [1,1] were analyzed by FACS. (H) Statistics of FACS data in (G). (n = 3). All data are displayed as the mean ± SD; \*\**p* < 0.01; statistical significance calculated using two-tailed Student’s t-test.

### Plasmids with identical overlapping sequences can construct Boolean AND logic circuits

We then subcloned sequence-overlapping ldsDNAs into the pMD-18T shuttle vector. A reporter assay showed that mammalian cells could conduct a Boolean AND gate response after being cotransfected with sequence-overlapping plasmids (Figure 9A). This result was further verified with pMD-18T vectors with overlapping GFP fragments (Figure 9D-F). In contrast to the case of sequence-overlapping ldsDNAs, there was a significant positive correlation between the length of the overlapping sequence and the output signal intensity under state [1,1] (Figure 9B). In addition, unlike sequence-overlapping ldsDNAs, downregulation of Rad51 could not decrease the state [1,1] signal from sequence-overlapping plasmids (Figure 9C).

**Figure 9.**
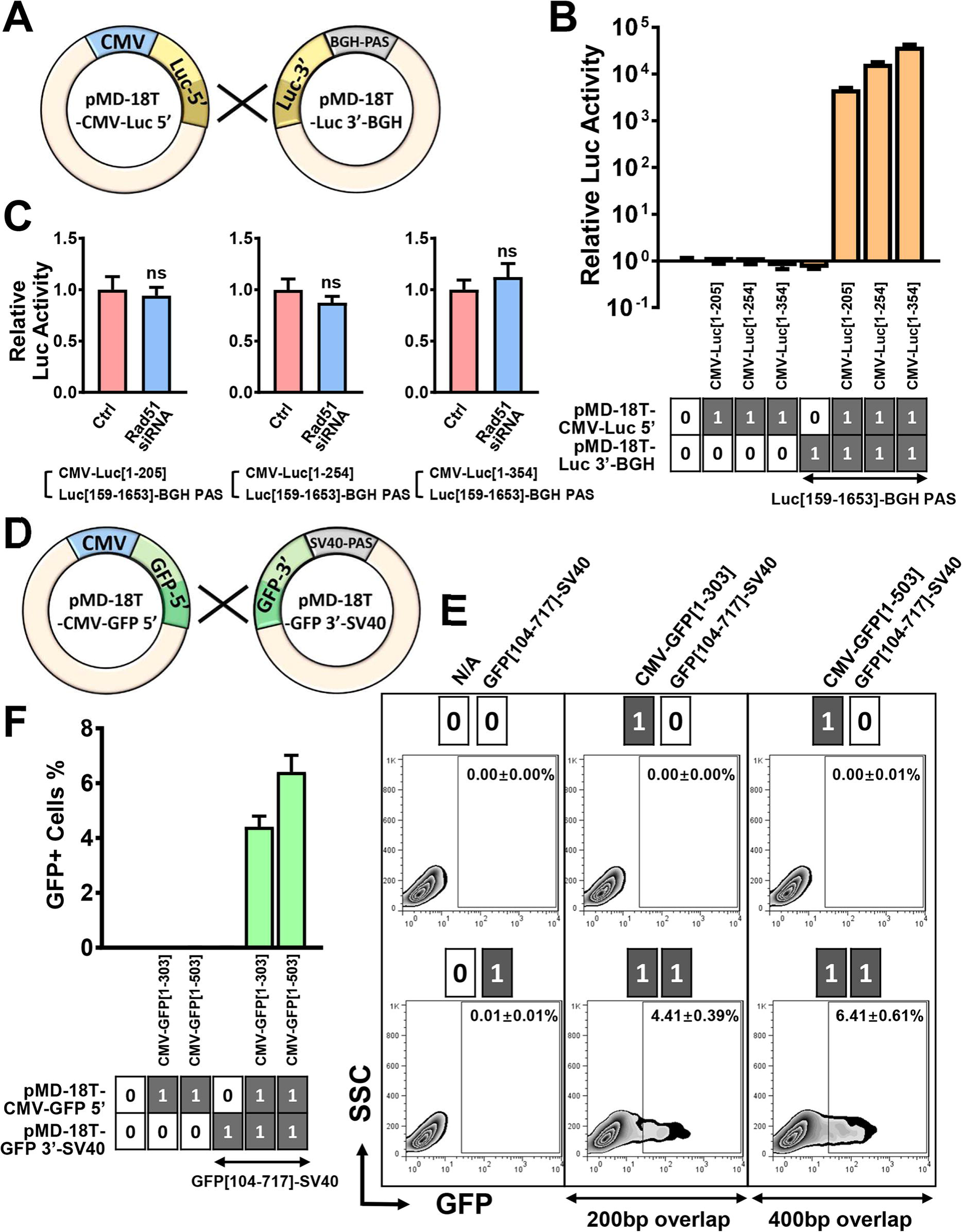
Boolean AND gate circuits by plasmids with identical overlapping sequences. (A) Schematic of plasmids with identical overlapping sequences: pMD-18T-CMV-Luc 5′ represents three plasmids in which CMV-Luc[1-205], CMV-Luc[1-254] and CMV-Luc[1-354] fragments were subcloned into pMD-18T; and pMD-18T-Luc 3′-BGH represents the pMD-18T plasmid with the Luc[159-1653]-BGH PAS fragment. (B) Different combinations of plasmids were transfected into HEK293T cells as indicated, and the firefly luciferase output signals under different states were measured and normalized to cotransfected Renilla signals. (n = 4). (C) Relative firefly luciferase activities of the indicated Boolean AND gate circuits (state [1,1]) in Rad51 knockdown cells. (n = 4). D) Schematic of plasmids with identical overlapping sequences: pMD-18T-CMV-GFP 5**’** represents two plasmids in which the CMV-GFP[1-303] and CMV-GFP[1-503] fragments were subcloned into pMD-18T, and pMD-18T-GFP 3**’**-SV40 represents the pMD-18T plasmid with the GFP[104-717]-SV40 PAS fragment. (E) Different combinations of plasmids were transfected into HEK293T cells as indicated, and the GFP-positive rates under different states were analyzed by FACS. (F) Statistics of FACS data in (E). (n = 3). All data are displayed as the mean ± SD; ns indicates not significantly different; statistical significance calculated using two-tailed Student’s t-test.

## Discussion

Here, by splitting the core gene expression cassette into signal-mute ldsDNA inputs, we present a proof-of-concept demonstration that ldsDNA could implement Boolean AND logic circuits in mammalian cells. Both luciferase and GFP expression could be achieved by ldsDNA-based Boolean AND logic circuits *in vitro*. Bioluminescence living imaging of firefly luciferase also demonstrated the feasibility of Boolean AND gate circuits *in vivo*. We also revealed that ldsDNA with terminal additional random nucleotide(s) could undergo end nucleotide deletion and generate in-frame protein via Boolean AND logic circuits. Moreover, ldsDNAs or plasmids with identical overlapping sequences could also form Boolean AND logic circuits in mammalian cells. This study provides novel biological parts and principles for synthetic genetic circuit design, which have application potential in future experimental and clinical studies.

Recently, several split-intein protein-splicing strategies have been developed to construct ‘bio-parts self-assembly’ AND logic circuits. Through split transcriptional regulatory elements, e.g., ZF-TFs, TALEs or dCas9, two- and three-input AND logic computation was implemented in cultured human cells [16-18]. All these systems are based on the expression of input intein fragment-transcriptional regulatory element domain fusion proteins, which perform transcriptional regulation. In contrast, our strategy does not require the prior expression of input proteins. The ldsDNA ligation reaction itself generates an intact full-length gene expression cassette, thereby inducing the expression of output signals. In split-intein strategies, given both the accessibility for intein trans-splicing reactions and the effects on protein folding, only limited split sites can be chosen. Our strategy has more freedom with regard to split sites. Either by splitting the promoter and CDS into different ldsDNAs or by splitting the CDS into different ldsDNAs, our circuit design achieves an excellent signal-to-background ratio: both luciferase assays and flow cytometry analysis showed ldsDNA-based circuit output signals thousands of times higher than single-input or background signals (Figure 1 and Figure 2).

In the current version, the two-input luciferase reporter signal level generated by our system was still lower than that generated by ldsDNA with an intact gene-expression cassette (Figure 1F and Figure 5). When the number of inputs comes to three, the output signal got even weaker (Figure 6). One possible factor limiting the efficiency is the intermediary process. As shown in our results, NHEJ machinery members exemplified by DNA-PKcs participate in output signal generation, implying that ldsDNA undergoes a rejoining process prior to the formation of the full-length productive gene-expression cassette. During the error-prone NHEJ process, nucleotide insertion or deletion occurs at DNA terminals [24, 25]. As shown in Figure 7, ldsDNAs with one or two additional terminal nucleotide(s) could also produce a GFP signal, indicating that the additional nucleotide(s) were deleted in the rejoining process to generate the in-frame GFP ORF. The addition of extra terminal nucleotide(s) leads to the reduction of both the GFP-positive rate and the mean fluorescence intensity. From another perspective, the in-frame ldsDNAs (without additional nucleotides) would also be subject to NHEJ end-processing, which may lead to the generation of alternative out-of-frame protein products. To avoid this, chemically modified ldsDNAs, which are resistant to nucleases, can be used for the construction of AND gate genetic circuits [26, 27]. Besides, we also took advantage of this characteristic and developed a novel methodology for high-content peptide library construction in mammalian cells, and deep sequencing revealed a high length-diversity of nucleotide sequences formed between ldsDNAs [28].

In this study, we also demonstrated that ldsDNAs or plasmids with identical overlapping sequences could implement Boolean AND gate circuits in mammalian cells. Sequence-overlapping ldsDNAs generate a stronger output signal than that of the ldsDNA end-joining strategy (Figure 1 and Figure 8). There was no significant correlation between the length of the ldsDNA overlapping sequence and the intensity of the output signals. Knockdown of Rad51, a crucial homologous recombination (HR) protein, could decrease the Boolean AND gate output signals generated by sequence-overlapping ldsDNAs, implying that the HR machinery would be involved in this process. Several studies have suggested a competition between HR and NHEJ response during double-strand DNA break repair, and inhibition of the NHEJ pathway was recently shown to improve the HR response [29-33]. Accordingly, small inhibitors of the NHEJ pathway may be utilized to improve output signals generated by sequence-overlapping ldsDNA-based genetic circuits [33]. In contrast to sequence-overlapping ldsDNAs, in sequence-overlapping plasmids, there was a positive correlation between the length of the overlapping sequence and the intensity of output signals, and Rad51 knockdown had no impact on Boolean AND gate output signals (Figure 9). The underlying mechanisms of these differences need to be further studied.

In summary, we developed several Boolean AND gate logic circuits utilizing ldsDNAs as re elements. This AND gate strategy was achieved both *in vitro* and *in vivo* in our study. We also showed that ldsDNA with terminal additional nucleotide(s) could undergo end chopping and output correct in-frame AND gate signals. Our study provides new biological parts and principles for synthetic biology and may have application potential in biomedical engineering fields.

## Materials and Methods

### Cell culture

The HEK293T cell line was maintained in Dulbecco’s modified Eagle’s medium (DMEM) (Thermo Fisher Scientific, Hudson, NH, USA) containing 10% fetal bovine serum (FBS) with 1% penicillin-streptomycin solution at 37°C with 5% CO_2_.

### Plasmids

Renilla luciferase control reporter vector (pRL-CMV, Promega, Madison, WI, USA) and GFP-expressing vector (pEGFP-C1, Clontech, Mountain View, CA, USA) were from lab storage. The firefly luciferase coding region was subcloned into the pcDNA3.0 vector (Invitrogen, Carlsbad, CA, USA) between the Hind-III and BamH-I restriction sites (namely, pcDNA3.0-firefly luciferase). The pMD-18T AT clone vector was purchased from TaKaRa (Beijing, China).

### ldsDNA synthesis

KOD DNA polymerase (Toyobo, Osaka, Japan) was employed to amplify Boolean AND gate ldsDNAs (PCR amplicons) using the pcDNA3.0-firefly luciferase or pEGFP-C1 plasmid as the template. To remove plasmid templates and free dNTPs, PCR products underwent agarose gel electrophoresis and gel purification (Universal DNA purification kit, Tiangen, Beijing, China). For animal experiments, ldsDNA was eluted in 0.9% NaCl solution. The concentration of PCR products was determined by the OD 260 absorption value.

### Transfection of Boolean AND gate genetic circuits

For the dual luciferase assay, HEK293T cells were seeded into 24-well plates the day before transfection (60%-70% confluency). Unless otherwise stated, 100 ng/well input amplicons or 400 ng/well input plasmids (for [1,1] states, 100 ng/well of each amplicon or 400 ng/well of each plasmid was used) and 10 ng/well pRL-CMV control vector were cotransfected into cells with Lipofectamine 2000 (Invitrogen) reagent following a standard protocol. Forty-eight hours after transfection, dual reporter activities were measured using the Dual Luciferase Reporter Assay System (Promega). Protein concentrations were determined with a Pierce BCA Protein Assay Kit (Thermo Fisher) according to the manufacturer’s instructions. Firefly luciferase activities (relative luminescence unit, RLU) were normalized either to Renilla luciferase activities (RLU) or protein concentrations (μg/μl).

For the GFP experiments, 100 ng/well input amplicons or 400 ng/well input plasmids were transfected into 24-well seeded HEK293T cells (for [1,1] states, 100 ng/well of each amplicon or 400 ng/well of each plasmid was used). Forty-eight hours after transfection, cells were harvested for FACS or western blot analyses.

### Immunofluorescence staining

For immunofluorescence staining, specific antibodies against γ-H2AX (05-636, Millipore, Billerica, MA, USA, 1:200 dilution) and 53BP1 (NB100-304, Novus, Centennial, CO, USA, 1:200 dilution) followed by Alexa Fluor 594 labeled-secondary antibodies (Invitrogen) were used for detection. Cells were counterstained with DAPI and mounted with anti-fade mounting medium, and photographs were taken with a confocal laser scanning microscope (Olympus, Tokyo, Japan).

### Flow cytometry analysis

HEK293T cells transfected with Boolean AND gate genetic circuits were rinsed with PBS and then digested with 0.25% trypsin-EDTA. Cells were resuspended in PBS supplemented with 2% FBS and then analyzed with BD FACSCalibur™ (BD, Bedford, MA, USA). Raw data were analyzed by FlowJo software.

### Western blots

Equal amounts of proteins were separated by SDS-PAGE and transferred onto PVDF membranes. After the membranes were probed with anti-GFP (ab1218, Abcam, Cambridge, MA, USA, 1:3000 dilution) or anti-β-actin antibody (sc-477778, Santa Cruz Biotechnology, Santa Cruz, CA, USA, 1:5000 dilution) and incubated with the appropriate secondary antibody, the antigen-antibody complex was visualized by chemiluminescent HRP substrate (Millipore).

### RNA interference and real-time PCR

siRNAs targeting DNA-PKcs (DNA-PKcs siRNA, CGU GUA UUA CAG AAG GAA ATT) [34] and Rad51 (Rad51 siRNA, GAG CUU GAC AAA CUA CUU CTT) [35] and scramble control siRNA (Ctrl, UUC UCC GAA CGU GUC ACG UTT) were synthesized by GenePharma (Shanghai, China). siRNAs were transfected using Lipofectamine 2000 reagent following the protocol. Total RNA was prepared 48 h after transfection and RT-qPCR was employed to analyze the interfering effects. cDNA was generated using the PrimeScript™ RT reagent Kit with gDNA Eraser (TaKaRa). Real-time PCR was performed using SYBR Premix Ex Taq™ II (Tli RNaseH Plus, TaKaRa). GAPDH mRNA levels were used for normalization. Forward (F) and reverse (R) primers used were as follows: GAPDH-F 5′- tcagtggtggacctgacctg -3′, GAPDH-R 5′- tgctgtagccaaattcgttg -3′; DNA-PKcs-F 5′- gctgatctcttaaagcgggc -3′, DNA-PKcs-R 5′- ctgtgtagcggatccaaacg -3′; Rad51-F 5′- tcacggttagagcagtgtgg-3′, Rad51-R 5′- ttagctccttctttggcgca -3′.

### Hydrodynamic injection of ldsDNA

All of the *in vivo* mouse experiments were approved by The Animal Ethical and Welfare Committee of Tianjin Medical University Cancer Institute and Hospital. Six- to eight-week-old syngeneic BALB/c female mice were purchased from the Beijing Vital River Laboratory Animal Technology Company (Beijing, China). 2.5 μg input ldsDNA (for [1,1] state, 2.5 μg of each input ldsDNA was used) was diluted with 1.6 ml normal saline at room temperature. Mice were anesthetized with 4% chloral hydrate and held immobile in a researcher’s hand, and then, the ldsDNA solution was injected into the retro-orbital sinus by a 26-gauge needle in 12 seconds.

### Bioluminescence imaging

BLI of live mice was performed on days 1, 2, 4, 7 and 14 after hydrodynamic injection of ldsDNA. BLI of firefly luciferase was performed using the IVIS Lumina II system (Xenogen, Alameda, CA, USA). After intraperitoneal injection of D-luciferin (Invitrogen, 150 μg/g body weight), each mouse was imaged for 1–5 min. Bioluminescence signals were quantified in units of maximum photons per second (photons/s).

BLI of mouse organs was performed one day after hydrodynamic injection of ldsDNA. Mice were injected with 150 μg/g D-luciferin intraperitoneally and were euthanized 10 min later. Different organs (heart, liver, spleen, lung, and kidneys) were collected immediately and placed into 6-well plates. BLI of organs was performed using the IVIS Lumina II system. Organs were imaged for 1–5 min. Bioluminescence signals were quantified in units of maximum photons per second (photons/s).

### Statistics

All data are presented as the mean ± SD unless stated otherwise. Differences were assessed by two-tailed Student’s t-test using GraphPad software. *p* < 0.05 was considered to be statistically significant.

## Abbreviations

ldsDNA: linear double-stranded
DNA NHEJ: non-homologous end joining
RLU: relative luminescence unit
BLI: bioluminescence imaging
CDS: coding sequence PAS
poly(A): signal DNA-PKcs
DNA-dependent protein kinase: catalytic subunit
HR: homologous recombinatio

## Acknowledgements

This work was supported by the National Natural Science Foundation of China [31870860, 31400673 to S.L., 81402407 to W.S.]; the Tianjin Research Program of Application Foundation and Advanced Technology [14JCQNJC09800 to S.L., 15JCQNJC11700 to W.S.]; and the Key Project of National Health and Family Planning Commission of Tianjin [2017057 to C.Z.].

## Author Contributions

S.L. conceived and supervised the study. S.L. and W.S. designed and performed the experiments. S.L. and W.S. analyzed the data. W.S. and S.L. prepared the figures. S.L. and W.S. wrote the paper. All authors discussed the results and reviewed the manuscript.

